# ETV2 regulates PARP-1 binding protein to induce ER stress-mediated cell death in tuberin-deficient cells

**DOI:** 10.1101/2021.11.08.467714

**Authors:** Shikshya Shrestha, Anthony Lamattina, Gustavo Pacheco-Rodriguez, Julie Ng, Xiaoli Liu, Abhijeet Sonawane, Jewel Imani, Weiliang Qiu, Kosmas Kosmas, Pierce Louis, Anne Hentschel, Wendy K. Steagall, Rieko Onishi, Helen Christou, Elizabeth P. Henske, Kimberly Glass, Mark A. Perrella, Joel Moss, Kelan Tantisira, Souheil El-Chemaly

## Abstract

Lymphangioleiomyomatosis (LAM) is a rare progressive disease, characterized by mutations in the tuberous sclerosis complex genes (*Tsc1 or Tsc2*), and hyperactivation of mechanistic target of rapamycin complex 1 (mTORC1). The effectiveness of mTORC1 inhibitors is limited by their lack of cytotoxic effects. Here, we report that E26 transformation specific (ETS) Variant Transcription Factor 2 (ETV2) is a critical regulator of Tsc2-deficient cell survival. Nuclear localization of ETV2 in Tsc2-deficient cells is mTORC1-independent and is enhanced by spleen tyrosine kinase (Syk) inhibition. In the nucleus, ETV2 transcriptionally regulates poly(ADP-ribose) polymerase 1 binding protein (PARPBP), a coregulator of transcription, mRNA and protein expression. Silencing of ETV2 or PARPBP in Tsc2-deficient cells induced ER-stress and increased cell death *in vitro* and *in vivo*. We also found ETV2 expression in human cells with loss of heterozygosity for *TSC2* lending support to the translational relevance of our findings. In conclusion, we report a novel signaling axis unique to Syk-inhibition is mTORC1-independent and promotes a cytocidal response in Tsc2-deficient cells, and therefore, maybe a potential alternative therapeutic target in LAM.

## Introduction

Lymphangioleiomyomatosis (LAM) is a rare, multisystem disease associated with smooth muscle-like “ LAM cells” over-proliferating in lungs, kidneys, and lymphatics. LAM cells have inactivating mutations in either tuberous sclerosis complex 1 or 2 (*TSC1* or *TSC2*) genes leading to hyperactivation of mechanistic/mammalian target of rapamycin complex 1 (mTORC1) signaling (Carsillo *et al*, 2000; Cheadle *et al*, 2000; Crino *et al*, 2006; Goncharova *et al*, 2002; Johnson *et al*, 2016; Rosset *et al*, 2017). A pivotal clinical trial has led to the approval of the allosteric mTORC1 inhibitor Sirolimus (rapamycin) by the Food and Drug Administrator for use in LAM (McCormack *et al*, 2011). However, due to the cytostatic nature, continuous treatment with rapamycin is required to maintain lung function stability (Yao *et al*, 2014). Therefore, there is a need for a better understanding of tuberin-deficient cell death mechanisms, which could potentially lead to novel therapies.

We have previously shown that there is increased expression and activation of spleen tyrosine kinase (Syk) in both Tsc2-deficient cells and LAM lung nodules (Cui *et al*, 2017). Similar to rapamycin treatment, R406, a Syk inhibitor (SykI) had antiproliferative effects in Tsc2-deficient cells *in vitro* and *in vivo* (Cui *et al*., 2017). Syk-dependent regulation of mTOR signaling has been previously documented in B-cell lymphoma (Fruchon *et al*, 2012; Leseux *et al*, 2006) and acute myeloid leukemia (Carnevale *et al*, 2013). We demonstrated that mTORC1-inhibition altered Syk expression and activity suggesting a feedback loop between the two pathways in Tsc2-deficient cells (Cui *et al*., 2017). Interestingly, R406 profiling has demonstrated that it has other kinase and non-kinase targets, including vascular endothelial growth factor receptor kinases (Fms related receptor tyrosine kinase 1 and kinase insert domain receptor), adenosine A3 receptor and cathepsins (Rolf *et al*, 2015). Here, we sought to investigate regulatory pathways of Syk-inhibition that are independent of its crosstalk with mTORC1 signaling in Tsc2-deficient cells and potentially identify target(s) of SykI that can have cytotoxic effects on Tsc2--deficient cells.

In this study, we performed an expression array of Syk- and mTORC1-inhibition to identify unique regulatory mechanism(s) between the two signaling pathways. We then utilized Passing Attributes between Networks for Data Assimilation (PANDA) (Glass *et al*, 2013), to reconstruct treatment-specific transcription factor regulatory networks in Tsc2*-*deficient cells. Using these networks, we identified E26 transformation specific (ETS) Variant Transcription Factor 2 (ETV2) that targeted unique set of genes in SykI network independent of mTORC1-inhibition and is a critical regulator of Tsc2-deficient cell survival.

ETV2 is a member of the ETS family of transcription factors, which plays a key role in the development of hematopoietic and endothelial lineages (Garry, 2016; Liu *et al*, 2015). ETV2 is expressed from embryonic days, E7.0 to E10.0 within the extraembryonic mesoderm, early progenitors, and newly formed endothelial cells, and *Etv2* deficiency is associated with a complete blockage in blood and endothelial cell formation (Garry, 2016; Oliver & Srinivasan, 2010). In zebrafish, *Etv2* expression is enriched in the posterior cardinal vein of developing embryos from where the lymphatic progenitors are derived. Furthermore, direct binding of ETV2 to the *Lyve1* and *Vegfr-3* promoter/enhancer results in the transcriptional regulation of those lymphatic markers (Davis *et al*, 2018). The role of ETV2 in lymphangiogenesis potentially has functional implications in LAM since there are lymphatic manifestations in LAM, including increased expression of VEGF-D in the serum of LAM patients (Glasgow *et al*, 2008; Seyama *et al*, 2006; Young *et al*, 2010). Studies have also demonstrated the potential lymphatic origin of LAM cells or Tsc2-deficient cells via expression of lymphatic markers, including PROX1, LYVE1, and VEGFR-3 in LAM cells (Davis *et al*, 2013) or Tsc2-deficient cells from angiomyolipoma (AML) (Yue *et al*, 2016). Recently, studies have demonstrated ETV2 mRNA expression and amplification in various tumor specimens, including glioblastoma and adrenocortical carcinoma (Li *et al*, 2018; Zhao *et al*, 2018). The activation of ETV2 in tumor-associated endothelial cells was shown to contribute to tumor angiogenesis (Kabir *et al*, 2018). There have been extensive studies exploring the function of ETV2 in the regulation of lineage development, and although some attention has been paid to exploring the role of ETV2 in human cancers, there are currently no data looking at ETV2 expression or function in tumor cells or cells that are of non-endothelial origin.

We demonstrated that ETV2 differentially regulates a unique set of genes in the SykI treatment network, including poly(ADP-ribose) polymerase 1 (PARP1) binding protein (PARPBP). PARPBP, also known as PARI or C12orf48, is a replisome associated protein that plays important roles during replication stress and DNA repair through its interaction with PARP-1, PCNA and RAD51 (Burkovics *et al*, 2016; Nicolae *et al*, 2019; Varisli, 2013). PARPBP overexpression is evident, and is associated with hyperproliferation, in pancreatic cancers, hepatocellular carcinoma, and myeloid leukemia (Nicolae *et al*., 2019; O’Connor *et al*, 2013; Piao *et al*, 2011; Yu *et al*, 2019). PARP-1 inhibitors are promising antineoplastic agents (Malyuchenko *et al*, 2015), and have been shown to selectively inhibit proliferation and induce apoptosis in Tsc2-deficient cells (Malyuchenko *et al*., 2015). However, the role of PARPBP in LAM or Tsc2-deficient cells and its regulation are largely unknown. Reporter assays showed that ETV2 transcriptionally upregulates PARPBP and silencing of ETV2 leads to decreased PARPBP mRNA and protein expression in Tsc2*-*deficient cells. Consequently, we observed increased endoplasmic reticulum (ER) stress and increased Tsc2-deficient cell death when ETV2 or PARPBP were silenced. Hence, targeting ETV2, with its potential for cytocidal cellular responses in Tsc2-deficient cells, might offer a therapeutic advantage in LAM over rapalogs alone, that primarily act as cytostatic drugs.

## Materials and Methods

### Cell culture

Tsc2-deficient Eker rat uterine leiomyoma (ELT3-V) cells were cultured using Dulbecco’s Modified Eagle’s medium (DMEM) containing 10% fetal bovine serum and incubated at 37°C in a humidified 5% CO_2_ atmosphere. For each experiment, cells were serum-starved overnight (16 hours) prior to treatment with vehicle control, DMSO (Sigma-Aldrich, St. Louis, MO), Syk-inhibitor (SykI) R406 (1µM; Selleckchem, Houston, TX), or rapamycin (20 nM; LC Laboratories, Boston, MA).

### GeneChip hybridization, and differential expression analyses

ELT3-V cells were treated with DMSO, rapamycin, or SykI for 24 hr (n = 4 biological replicates per group). RNA was extracted with the RNeasy Mini Kit (Qiagen, Hilden, Germany). All subsequent sample preparation and GeneChip (Rat Gene 2.0 ST arrays) processing for microarray analysis were performed by the Boston University Microarray and Sequencing Resource Core facility using an input of 1 µg of total RNA from each sample. Analysis was performed using the Bioconductor software suite (version 2.12) (Gentleman *et al*, 2004). Chip definition file (CDF) (Dai *et al*, 2005) was processed using a robust multi-array average (RMA) algorithm (Irizarry *et al*, 2003) available in the *affy* package (version 1.36.1) (Gautier *et al*, 2004). The log_2_ scale data from RMA was used in statistical testing.

Differential expression analysis was performed and Benjamini-Hochberg false discovery rate (FDR) (Benjamini & Hochberg, 1995) was implemented for both ANOVAs and *t*-tests to generate corrected *p* values (*q* values). A filtered gene list was generated for expression changes of greater than 2.0-fold and one-way ANOVA FDR *q* < 0.01 and, furthermore, divided into four distinct clusters based on expression pattern in the three conditions to generate a heat-map. Principal Component Analysis (PCA) was performed by normalizing gene expression values across all samples to a mean of 0 and a standard deviation of 1.0 with the *prcomp* R function. The three treatment conditions were separated with the first principal component (PC1), and replicates in each condition were separated with the second PC2.

### Pathway analyses

GO term enrichment analysis of clustered gene groups was performed using DAVID (Huang da *et al*, 2009a, b) with default settings. HomoloGene (version 68) (Coordinators, 2013) was utilized to identify human homologs of the rat genes in the array. The R environment (version 3.4.3) was used for all microarray analyses. Kyoto Encyclopedia of Genes and Genomes Pathway analysis (KEGG Pathway analysis) was used to perform subsequent bioinformatics analysis of all the genes identified in microarray, irrespective of the clusters. For both GO term and KEGG analysis pathways were selected with a *P*-value <0.05 and gene count >2.

### Transcription factor and target network construction

Passing attributes between networks for data assimilation (PANDA) analysis was utilized to construct gene regulatory networks for each of three treatment drugs, Syk, Rapa, and DMSO using all genes in the microarray dataset, as previously described (Glass *et al*., 2013). An initial map of transcription factors to genes was created by scanning the rn6 genome for 620 Cis-BP Rattus norvegicus motifs provided with the MEME suite (Bailey *et al*, 2009) using the Finding Motif Occurrences (FIMO) program (Grant *et al*, 2011). Statsitically significant (p<1e-4) hits within the promoter region, defined as a the [-750,+250] bp region around the transcriptional start site of RefSeq annotated genes, were retained. The initial mapping included 2,245,143 edges (17,177 genes, 616 TF). There were 14,890 genes and 616 TF common to both expression data and the motif mapping. Potential inferred regulatory relationships were determined by using PANDA to integrate the motif mapping and gene expression data and assign a weight (z-score) to each edge that connects a TF to its target gene. Top 10,000 (TF, gene) pairs with the largest absolute differences of edge weights in three different pairwise comparisons were generated: Pair 1. E(SykI)-E(DMSO), Pair 2. E(rapamycin)-E(DMSO), and Pair 3. E(rapamycin)-E(SykI). Additionally, edge weight differences for Pair 3 were plotted as scatter plots for visualization using Cytoscape. Transcription factors involved in top edges for each pairwise comparison were identified from the network of differential regulation.

### Real-Time Polymerase Chain Reaction (RT-PCR)

One µg of total RNA extracted from ELT3-V cells with the RNeasy Mini Kit (Qiagen, Germantown, MD) was reverse-transcribed into cDNA using amfiRivert cDNA Synthesis Master Mix (GenDEPOT, Barker, TX). Quantitative real-time polymerase chain reaction (qPCR) was performed using iTaq Universal SYBR Green qPCR Master Mix (Biorad, Hercules, CA), according to the manufacturer’s protocol. Supplementary Table 1 provides the primer sequences used in qPCR analyses.

### Cellular Fractionation

Treated ELT3-V cells were washed with cold PBS and collected by scraping. Cellular fractionation was performed using a CelLytic™ NuCLEAR™ Extraction Kit (Sigma-Aldrich, St. Louis, MO), according to the manufacturer’s instructions. A portion of cell pellets was used for total protein isolation using RIPA buffer (Thermo Fisher Scientific, Waltham, MA). Both nuclear and total protein isolation buffers were supplemented with protease and phosphatase inhibitors (Invitrogen, Carlsbad, CA). Equal fractions of lysates for both nuclear and cytoplasmic fractions were subjected to immunoblotting.

### Immunoblot

Equal amounts or fractions of indicated protein lysates were loaded onto NuPage 4-12% Bis-Tris Protein Gels (Invitrogen, Waltham, MA), and then subsequently immunoblotted with the primary antibodies listed in Supplementary Table 2. Each protein of interest was then detected with HRP-conjugated goat anti-rabbit or anti-mouse IgG antibody (1:2000; Invitrogen), and visualized using SuperSignal West Pico PLUS Chemiluminescent Substrate or SuperSignal™ West Femto Maximum Sensitivity Substrate (Thermo Fisher Scientific, Waltham, MA).

### RNA interference

Predesigned MISSION siRNA targeting rat Etv2 and rat Parpbp, as well as MISSION siRNA Universal Negative control, were purchased from Sigma-Aldrich. ELT3-V cells were transfected with Etv2 siRNA (10 uM), Parpbp siRNA (5 uM), or control siRNA (SCR) using Lipofectamine RNAiMAX reagent (Thermo Fisher Scientific) and OPTI-MEM (Thermo Fisher Scientific). Sequences for siRNA used are listed in Supplementary Table 3.

### Arsenite-induced stress granule formation

ELT3-V cells were plated in 4-well chamber tissue culture slides (Corning, Corning, NY) and transfected with SCR or Etv2 siRNA. Forty-eight hours post-transfection, cells were treated with 0.5 mM sodium arsenite (NaAsO_2_) (MilliporeSigma, Burlington, MA) for 40 min. The cells were then fixed with 4% paraformaldehyde and processed for immunofluorescence analysis using GAP SH3 Binding Protein 1 (G3BP1) antibody (green; Abcam, Cambridge, MA) and fluorophore-conjugated secondary antibody. Nuclei were visualized with 4’,6-diamidino-2-phenylindole (DAPI, blue; Sigma) staining. Images were captured with a FluoView FV-10i Olympus Laser Point Scanning Confocal Microscope using a 60X objective (Olympus, Tokyo, Japan).

### Luciferase reporter assay

The *Parpbp* promoter fragments with (386 bp) or without (150 bp) one putative ETV2 binding site (Wei *et al*, 2010) were cloned into pGL3_Basic vector (Cat# E1751, Promega, Madison, WI). Primers Forward: taagcagagctcgtcggagggcgagcgaggcg and Reverse: tgcttaaagcttttaccacgatgccgctggagg were used to clone −311 to +75 bp and primers Forward: taagcagagctcggcgcggaacaagcgtagtagtcag and Reverse: tgcttaaagcttttaccacgatgccgctggagg were used to clone −75 to +75 bp promoter fragments. Reverse transfection was carried out in a total of 15,000 ELT3-V cells transfected with SCR and Etv2 siRNA for 24 hours using X-tremeGENE HP DNA transfection reagent (Sigma). Cells were transfected with 10 ng of pRL_CMV vector (internal control, Cat# E2261, Promega) and 140 ng of promoter constructs. Luciferase activity was tested 24 hours after plasmid transfections using the Dual-Glo Luciferase Assay System kit (Cat# E2920, Promega) as per the manufacturer’s protocol. Each transfection was carried out in triplicates in 4 independent experiments. Biotek Synergy HT microplate reader (BioTek, Winooski, VT) with Biotek Gen5.1.1 microplate data collection software was used for luciferase luminescence detection.

### Animal studies

All animal experimental procedures were performed according to protocols approved by the Institutional Animal Care and Use Committee at Brigham and Women’s Hospital. ELT3-V cells stably transduced with pCMV-Luciferase (ELT3-V-Luciferase) were transfected with Scr siRNA and Etv2 siRNA. 24hr post-transfection, 1 × 10^6^ cells were injected into female immunodeficient C.B17 SCID mice (Taconic, Hudson, NY) via lateral tail-vein injections. Before imaging, mice were injected with RediJect D-luciferin bioluminescent substrate (Catalog#770504; Perkin-Elmer, Waltham, MA). Bioluminescent signals were recorded at 4 hr, 24 hr, and 48 hr using an In-Vivo Xtreme imaging system (Bruker, Billerica, MA). Net luminescence intensity in the chest area was assessed using Bruker molecular imaging (MI) software (v7). Murine blood was collected at the end of the experiment by aortic puncture and red blood cells were lysed using an ammonium-chloride-potassium (ACK) lysing buffer (Quality Biological, Gaithersburg, MD). Additionally, murine lung tissues were harvested, minced, and enzymatically digested (300 units/ml Collagenase 4, Worthington Dorchester, MA). DNA was extracted from blood and mouse lung tissues using a Blood and Tissue DNA kit (Qiagen, Germantown, MD). Rat and mouse DNA were quantified by RT-qPCR using rat- and mouse-specific primers included in Supplementary Table 1 (Walker *et al*, 2004; Yu *et al*, 2009).

### Analysis of available single-cell RNA-Seq data

scRNA seq-data for 4 LAM samples were downloaded from Gene Expression Omnibus (GSE135851) (Guo *et al*, 2020). Previous analysis of LAM sample 2 did not detect LAMCORE positive cells and therefore was excluded from our analysis. The data from LAM samples 1, 3, and 4 were analyzed using Seurat v3 (Butler et al., 2019). For the 3 LAM samples, genes expressed in at least two cells were selected. Furthermore, for LAM samples 1 and 3 specifically, cells with at least 500 expressed genes (nFeature_RNA >500) with less than 10% of genes mapping to the mitochondria (percent.mt < 10) were selected. For LAM sample 4, nuclei with at least 300 expressed genes and with less than 10% of genes mapping to the mitochondria were included for analysis. The three Seurat objects were merged into a single object and preprocessed using the outline previously described (Butler et al., 2019). The samples were then integrated using the R package Harmony (Korsunsky et al., 2018) followed by imputation of the data using the R package Alra (Linderman et al., 2018). Clusters were then generated using Seurat and labeled with the R package ClustifyR (Fu R, Gillen AE, Sheridan RM, et al. 2020) using the data published by Habermann et al. 2019 as a reference matrix. 121 LAMCORE cells were identified based on positive (>0) expression of LAM markers PMEL, FIGF, ACTA2, and VEGF-D (Guo *et al*., 2020); 100 of these cells clustered together in the LAMCORE cluster. 96 ETV2 positive cells within the LAMCORE cells were identified by positive (0>) expression of ETV2.

### Isolation of cells with loss of heterozygosity for *TSC2*

Cells isolated from explanted lungs of patients undergoing lung transplantation were cultured in mesenchymal stem cell media (MSCGM) as described previously (Pacheco-Rodriguez *et al*, 2007). Cultured cells (∼1×10^6^) were reacted with 20 µl antibodies of each anti-CD44v6-FITC and anti-CD44-PE (Supplementary Table 2) following trypsinization. Cells underwent a 30-minute incubation at room temperature. Then, 3 ml of PBS were added and the tubes were centrifuged for 10 min at ∼250 × *g*. The cell pellet was then suspended in 500 µl PBS. Cell mixtures were placed on ice until sorted using a BD FACSaria 2 (Becton Dickinson, Franklin Lakes, NJ). Four subpopulations of cells were obtained in PBS and RNA later (Sigma, St. Louis, MO). We previously showed that cells sorted expressing CD44 and CD44v6 were more likely to have loss of heterozygosity (LOH) for *TSC2*. To identify circulating LAM cells, we took 60 ml of heparinized fresh blood to isolate cells following a density gradient. For that purpose, we used CD235a-PE and CD45-FITC antibodies (Supplementary Table 2). Cells with *TSC2* LOH are most likely to be found in cells expressing CD235a. *TSC2* LOH was determined using five microsatellite markers (D16S521, D16S3024, D16S3395, Kg8, and D16S291) on chromosome 16, as previously described (Steagall *et al*, 2013).

Total RNA was isolated from cells in RNALater, cDNA synthesized, and qPCR performed, according to the manufacturer’s protocol. Primer sequences used in qPCR analyses were provided in Supplementary Table 1.. PCR products were subjected to agarose gel (2%) electrophoresis to visualize the amplified ETV2 qPCR products.

### Statistical analysis

For all experiments, at least three independent experiments were conducted, and data are presented as mean ± standard error of the mean (SEM). Statistical analyses of all endpoints were performed using one-way ANOVA, followed by a Tukey *post hoc* test or two-tailed Student’s t-test. *p* < 0.05 was considered statistically significant. Analyses were performed using GraphPad Prism 8.3 (GraphPad Software, La Jolla, CA).

## Results

### Gene expression profiling in rapamycin-versus SykI-treated Tsc2-deficient cells

To discern the effects on gene expression of mTORC1 and Syk inhibition in Tsc2-deficient cells, we ran Rat Gene 2.0 ST microarrays with ELT3-V cells treated with DMSO, SykI, or rapamycin for 24 hours. A Heatmap generated using the top 485 most differentially expressed genes in each treatment condition compared to control (DMSO) showed four distinct clusters (Figure 1A). We validated the gene expression data for the top 8 differentially expressed genes from each cluster using RT-qPCR (Figure S1A-D). PCA plot, based on PC1, revealed DMSO-treated replicates clustered very distinctly from both SykI- and rapamycin-treated replicates, suggesting considerable changes in gene expression in the treatment groups compared to DMSO (Figure 1B). In contrast, only a minute separation in the clusters was observed between the two treatment groups suggesting largely concordant changes in gene expression (Figure 1B).

**Figure 1.**
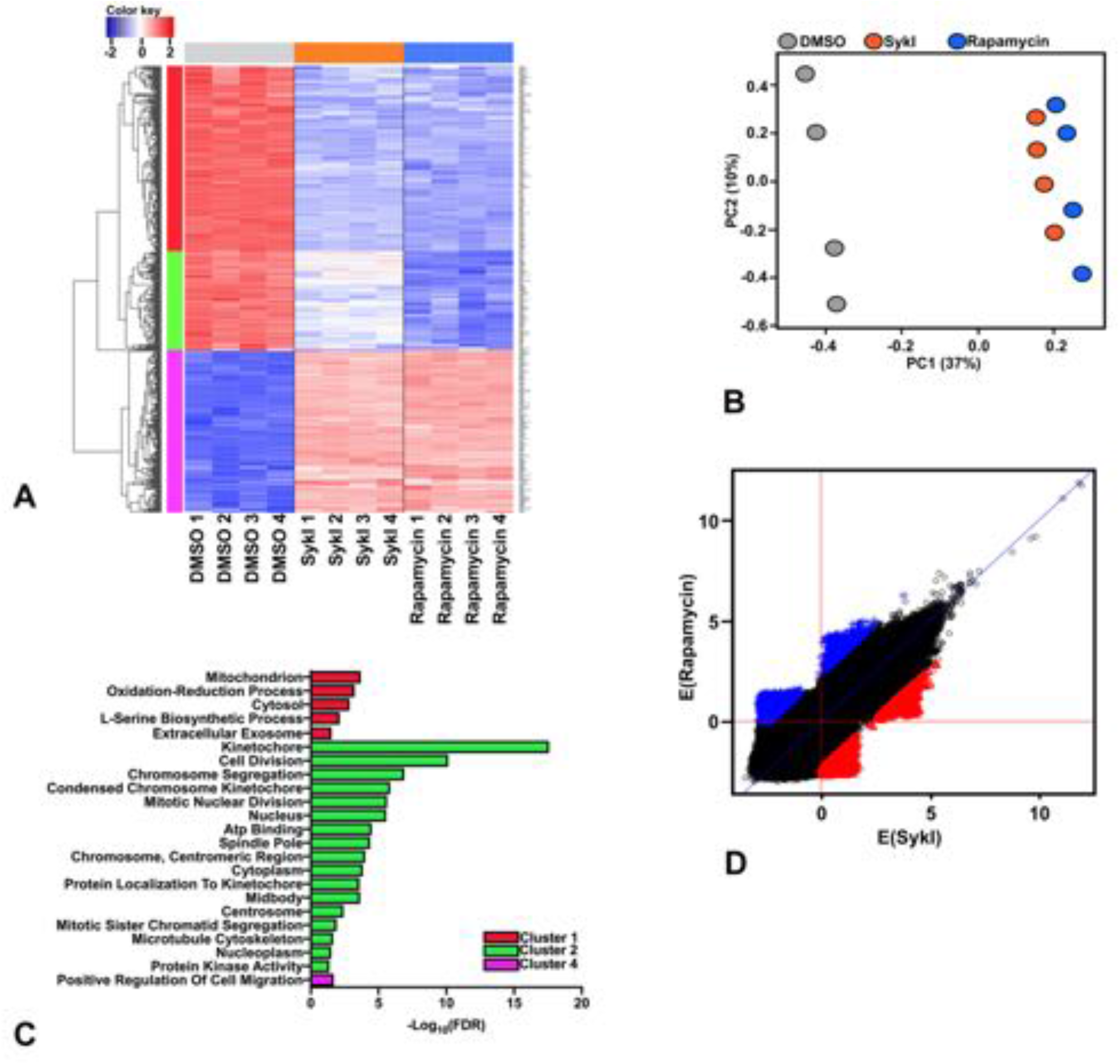
Gene expression profiling in rapamycin-vs. SykI-treated ELT3-V cells. A) *Tsc2*-deficient Eker rat (ELT3-V) cells were treated with SykI (R406; 1 uM), rapamycin (20 nM), or DMSO for 24 hours. Data were obtained using Affymetrix Rat Gene 2.0 ST GeneChip. The diagram represents 485 differentially expressed probes (*q* < 0.01) with at least two-fold change between conditions. Each *column* represents a single experiment, and each *row* a single gene. The colors are scaled so that red and blue indicate z-scores of ≥ 2 or ≤ −2, respectively, and white indicates a z-score of 0 (row-wise mean). Color blocks on top, gray, orange, and blue represent different treatment groups, DMSO, SykI, and rapamycin, respectively. Color blocks on the left, red, green, blue, and pink represent 4 different clusters, Clusters 1 - 4. B) Principal Component Analysis (PCA) of samples treated with DMSO, rapamycin, or SykI for 24hr. PCA was performed across all probes on the microarray and a bi-plot of PC1 against PC2 was plotted to show clustering of different treatment groups and condition replicates. C) GO enrichment analysis for significantly upregulated biological processes using genes in each cluster in the heatmap (A). GO-terms for Cluster 1, 2, and 4 are presented in red, green, and pink, respectively. D) Scatter plot for the SykI and rapamycin networks predicted by PANDA for the top 10,000 edges (TF, gene) in each of the networks models, rapamycin-treatment, and SykI-treatment models. Each point in the graph represents a difference in edge weight connecting a transcription factor to a target gene between two networks. Red points represent the 10,000 (TF, gene) pairs having largest weight in SykI network and blue points represent (TF, gene) edges having the largest weight in rapamycin network.

The top cluster (Cluster 1, red) in the heatmap included 201 genes, which were similarly downregulated in both SykI- and rapamycin-treated cells compared to DMSO-treated cells (Figure 1A). Gene ontology (GO) enrichment analysis on these genes showed upregulation in biological processes that include oxidation-reduction and L-Serine biosynthesis (Figure 1C). The second cluster (Cluster 2, green) included 107 genes that were downregulated in both treatment groups compared to DMSO but demonstrated greater downregulation with rapamycin than with SykI (Figure 1A). These genes corresponded to 17 significantly upregulated biological processes, most of which were associated with the cell cycle (Figure 1C). Only one gene (*Mospd1*), was downregulated by SykI treatment, but not rapamycin, comprising the third cluster (Cluster 3, blue). RT-qPCR validation, however, showed downregulation of *Mospd* in both SykI and rapamycin treatment groups compared to DMSO (Figure S1C). The fourth cluster (Cluster 4, pink) included 176 genes that were upregulated in both treatment groups compared to DMSO, and corresponded to a significant upregulation of biological processes associated with cellular migration (Figure 1A and 1C).

Overall, differential expression data from the microarray indicated that SykI and rapamycin treatments similarly impacted gene expression in Tsc2-deficient cells. A subset of genes, comprising Cluster 2, were observed to be more downregulated with mTORC1 inhibition compared to Syk inhibition.

### Network analysis reveals putative SykI-specific regulation driven by the transcription factor Etv2

To further understand potential gene regulation differences between SykI and rapamycin treatments, we used PANDA analysis to integrate information from treatment-specific gene expression and transcription factor binding motifs and construct transcriptional regulatory networks for the three treatment groups using pairwise comparisons between two groups at a time: SykI vs. DMSO, rapamycin vs. DMSO, and SykI vs. rapamycin. Edge weight differences for the top 10,000 (TF, gene) pairs with the largest absolute differences of edge weights between 2 networks SykI and rapamycin were plotted (Figure 1D). Consistent with the finding from differential gene expression and pathway analyses, a high correlation between the edge weights for the two treatment groups were observed; however, we also found regulatory edges that were more strongly supported by either SykI (red) or rapamycin (blue) treatment. Based on the top 10,000 (TF, gene) pairs, SykI and rapamycin networks consisted of 54 unique TFs in either of the two networks, and most TFs belonged to the members of the E26 transformation specific (Ets) family of TFs (Table 1). *Ets* variant 2 (Etv2) was among the top 20 TFs regulating the highest numbers of genes in each network. Since evidence suggested a potential lymphatic endothelial origin of LAM cells (Davis *et al*., 2013; Yue *et al*., 2016) and that ETV2 is critical for lymphangiogenesis ((Davis *et al*., 2018), we further investigated the role of ETV2.

**Table 1.**
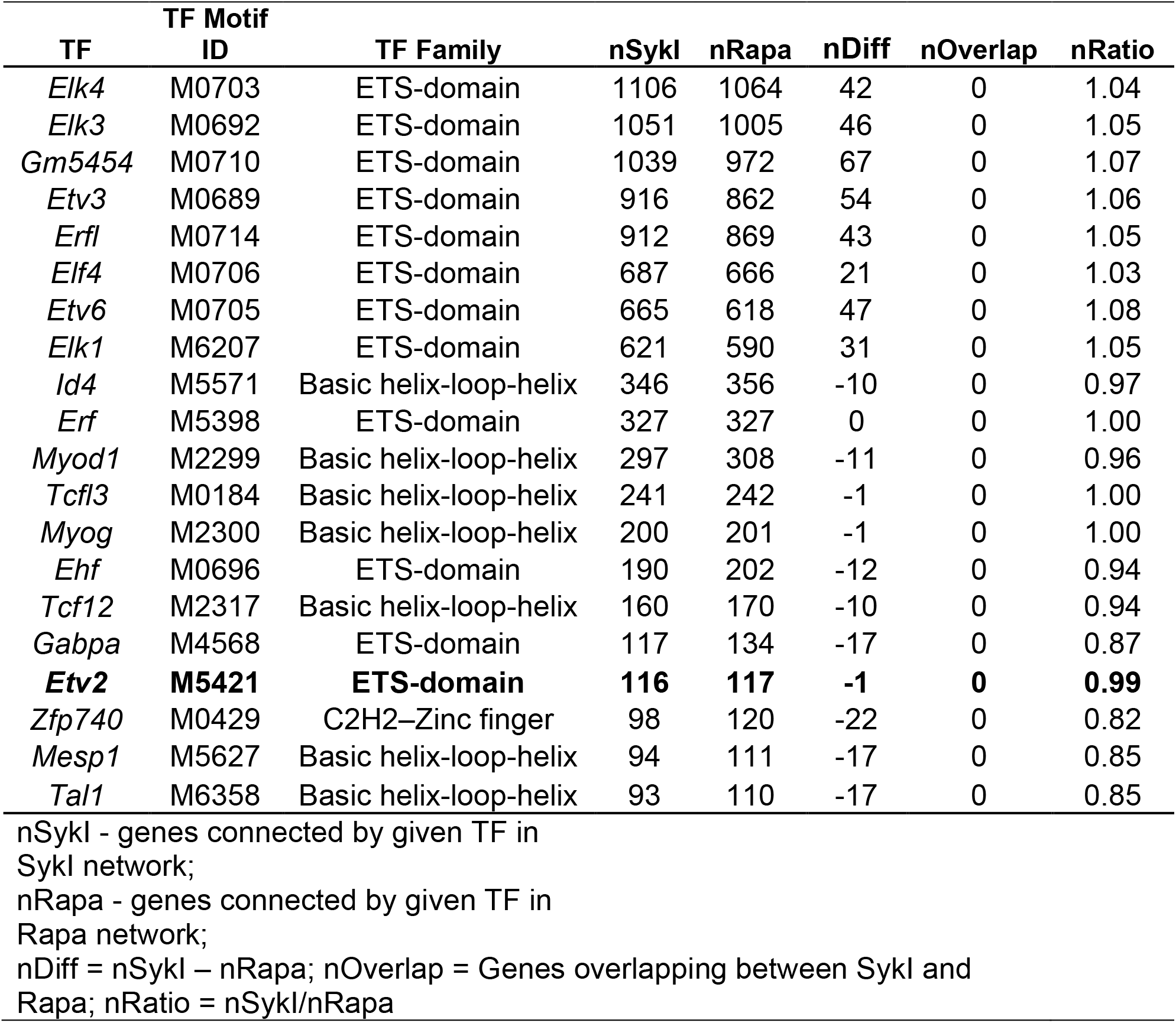
The TF with highest nDiff in SykI and Rapa networks identified using top 10,000 (TF, genes) pairs.

### SykI, but not Rapamycin treatment induced ETV2 translocation into cell nuclei

To understand a potential role for ETV2 in mediating SykI-dependent function in Tsc2-deficient cells, we investigated *Etv2* gene and ETV2 protein expression in Tsc2-deficient ELT3-V cells treated with DMSO, SykI, or rapamycin for 24 hours. No significant differences in both total mRNA and total protein expression were observed with either SykI or rapamycin treatment compared to DMSO (Figure 2A-C). SYK has previously been shown to regulate the nuclear translocation of transcription factors, including nuclear factor (erythroid-derived 2)-like 2 (Nrf2) and Ikaros (Park *et al*, 2018; Uckun *et al*, 2012). Therefore, we also examined the effect of SykI and rapamycin treatment on ETV2 localization. Western blot analysis of ELT3-V cell fractions showed significantly increased nuclear ETV2 protein levels in SykI-treated cells compared to DMSO-treated cells (Figure 2D-E), and importantly treatment with rapamycin did not result in ETV2 nuclear accumulation. Additionally, immunofluorescence corroborated the finding and demonstrated that SykI-treated ELT3-V cells featured higher nuclear levels of ETV2 than both DMSO- and rapamycin-treated cells after 24 hours (Figure 2F). These data collectively suggest that Syk-inhibition, but not mTORC1-inhibition drives ETV2 nuclear translocation.

**Figure 2.**
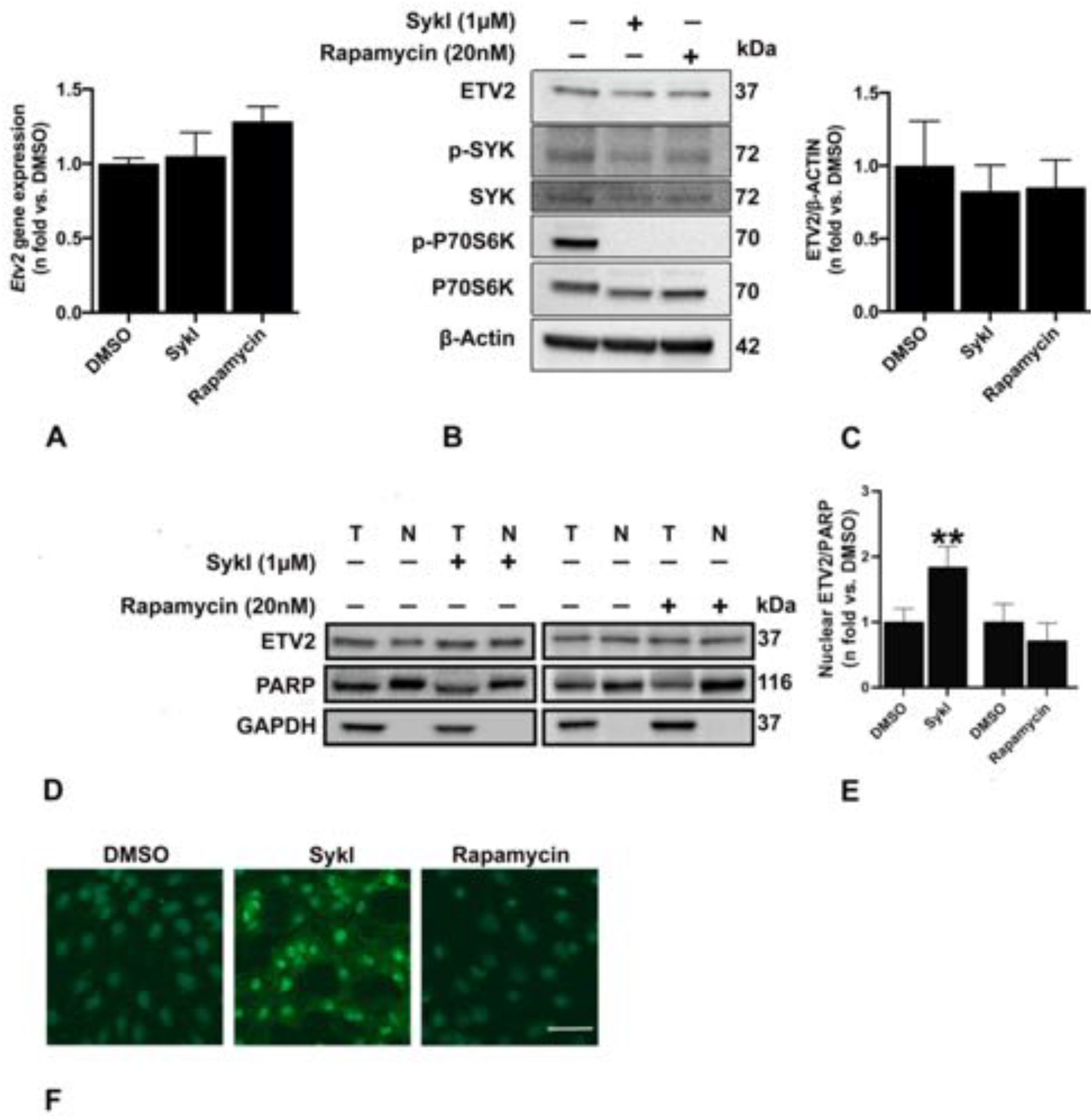
Effect of SykI and rapamycin treatment on Etv2 expression and localization. A) Real-time qPCR analysis of rat Etv2 mRNA in *Tsc2*-deficient ELT3-V cells treated with SykI, or rapamycin for 24hr compared to DMSO-treated cells. Data represent mean ± SEM of three independent experiments. B-C) Immunoblot analysis of equal amount of rat ETV2 protein from ELT3-V cells treated with DMSO, SykI, or rapamycin for 24hr. Antibodies against rat ETV2, p-SYK, SYK, p-P70S6 kinase, and P70S6 kinas*e* were used, and β-Actin was used as a loading control. Band intensities of ETV2 were assessed, and ratios of ETV2/β*-*Actin were calculated for each treatment group. Results were expressed relative to DMSO. Data are means ± SEM of at least three independent experiments (ETV2/β*-*Actin: P > 0.05; Student’s t-test). D-E) ELT3-V cells were treated with DMSO, SykI, or rapamycin as in (B). Samples from equal fractions of total (T) and nuclear (N) protein lysates were separated by SDS-PAGE and transferred to PVDF membranes, which were incubated with antibodies against ETV2, PARP (nuclear marker), and GAPDH (cytoplasmic marker). Band intensities of ETV2 were analyzed, and ratios of ETV2/PARP and ETV2/GAPDH were calculated for each group. Results were expressed relative to DMSO. Data are means ± SEM of at least three independent experiments (ETV2/PARP: *P < 0.05, **P < 0.01; Student’s t-test). F) Merged images of ETV2 (*green*) and 4’,6-diamidino-2-phenylindole (DAPI; *blue*) fluorescence of representative ELT3-V cells treated as indicated with DMSO, SykI, or rapamycin. Scale bar equals 150µm. **Figure 2-Source data 1** Table showing data from experiments plotted in Figure 2A, 2B and 2E. **Figure 2-Source data 2** Original, unedited western blot image files and original images with each band labeled for Figure 2C **Figure 2-Source data 3** Original, unedited western blot images and original images with each band labeled for Figure 2D

### ETV2 silencing induces ER stress and leads to Tsc2-deficient cell death

In addition to its role in endothelial and lymphatic lineage, ETV2’s role as an important regulator of endothelial cell survival, cell cycle, and proliferation has been previously studied (Abedin *et al*, 2014; Singh *et al*, 2019). To assess the effect of ETV2 alteration on Tsc2-deficient cells, we genetically silenced Etv2 in ELT3-V cells (Figure 3A-C). Silencing ETV2 lead to a significant increase in Annexin V cells, indicating increased cell apoptosis/death with reduced ETV2 expression (Figure 3D-F). We then sought to determine whether the ETV2 silencing-mediated cell death is related to endoplasmic reticulum (ER) stress since Tsc2-deficient^-^ cells are known to be sensitive to ER stress (Kang *et al*, 2011; Ozcan *et al*, 2008). As shown in Figure 3, expression of ER stress markers CHOP (Figure 3G-H) and phosphorylated EIF-α (Figure 3G and 3I) were markedly elevated in ETV2-silenced ELT3-V cells, suggesting increased ER stress. Importantly, the ER stress response was associated with increased cell death as demonstrated by the increase in cleaved PARP (Figure 3G and 3J).

**Figure 3.**
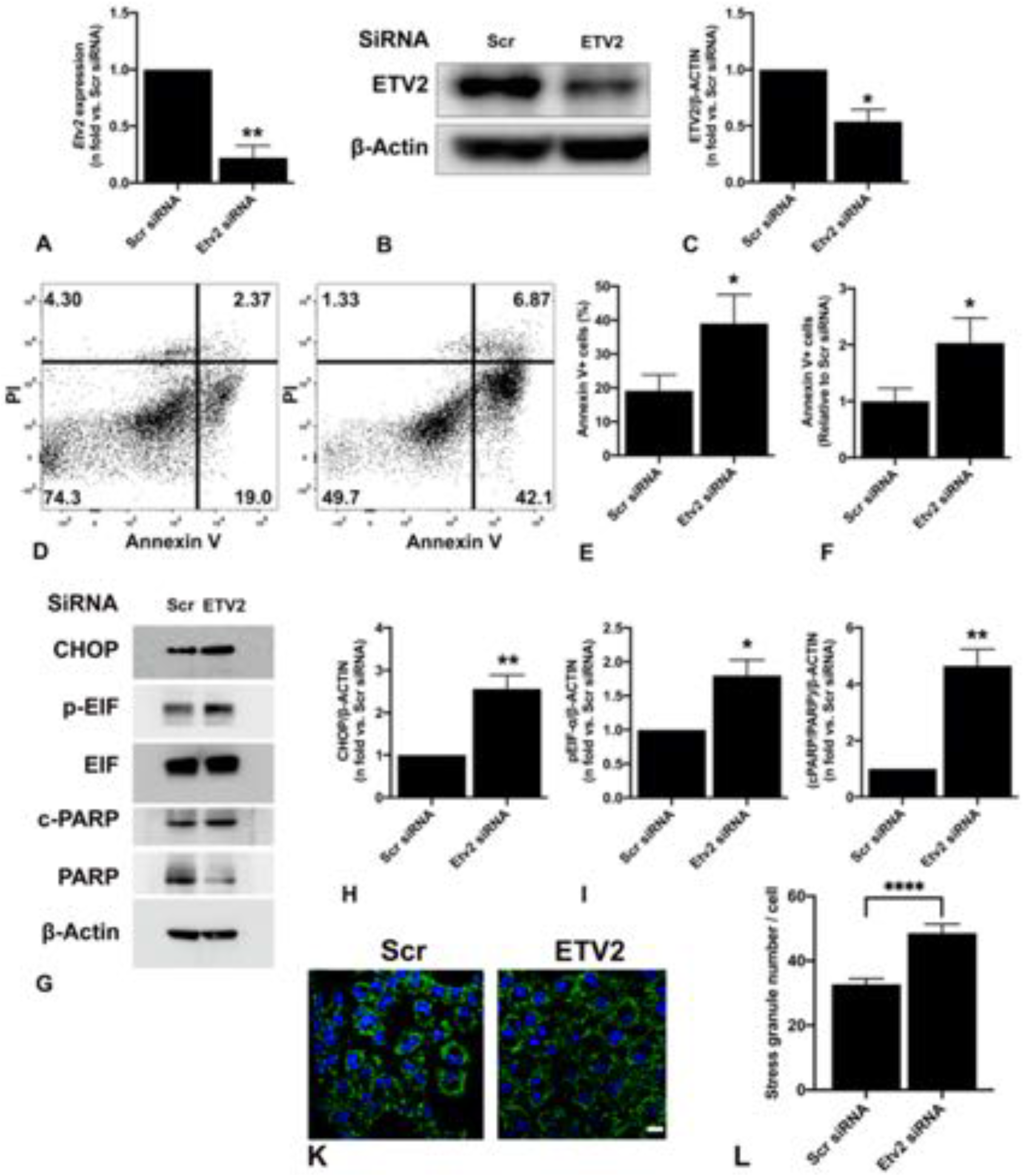
Silencing ETV2 induces ER stress and cell death in Tsc2-deficient cells. *Tsc2*-deficient cells (ELT3-V) were transfected with either SCR or Etv2 siRNA for 48h. A)Real-time qPCR analysis of rat Etv2 mRNA. The histogram represents fold change in mRNA expression compared to SCR control. Data are means ± SEM of at least three independent experiments (**P < 0.01; Student’s t-test). B-C) Equal amounts of protein from whole-cell lysates were analyzed by Western blotting with antibodies against ETV2 and β-actin. The ratio of ETV2 to β-actin density was expressed as the fold-change relative to Scr siRNA. Data are means ± SEM of at least three independent experiments (*P < 0.05; Student’s t-test). D-F) PI and FITC AnnexinV apoptosis assay and flow cytometry were used to measure cell apoptosis induced by ETV2 silencing in ELT3-V cells. Representative flow cytometry plots for SCR and Etv2 siRNA are presented. The total percent of Annexin V positive (+) cells in each group were quantified, and Annexin V+ cells in Etv2 siRNA-transfected cells relative to Scr siRNA-transfected cells calculated from 3 independent experiments (*P < 0.05; Student’s t-test). G-J) Equal amounts of proteins were separated by electrophoresis and transferred to PVDF membranes, which were reacted with antibodies against CHOP, pEIF-α, total EIF-α, cleaved (c)-PARP, and total PARP.β-Actin was used as a loading control. Ratios of CHOP (H), pEIF-α/EIF-α (I), and c-PARP/PARP(J) to β-Actin were expressed as fold change to control siRNA (Scr siRNA). Data represent means ± SEM of at least three independent experiments (*P < 0.05, **P < 0.01; Student’s t-test). K-L) ELT3-V cells were treated with 0.5 mM NaAsO_2_ (Arsenite) for 40 min. Cells were stained for G3BP1 (green) to detect stress granules and DAPI (blue) was used to visualize the nuclei. A total of 221 cells per siRNA group were imaged and quantified with CellProfiler. Scale bar, 20 µm, ****p<0.0001 **Figure 3-Source data 1** Table showing data from experiments plotted in Figure 3A, 3C, 3E, 3F, 3H, 3I, 3J and 3L. **Figure 3-Source data 2** Original, unedited western blot images and original images with each band labeled for Figure 3B **Figure 3-Source data 3** Original, unedited western blot image files and original images with each band labeled for Figure 3G

To further confirm the ETV2-mediated induction of ER stress, we examined the role of ETV2 in the regulation of stress granule (SG) dynamics. We evaluated oxidant-induced stress granule formation in ELT3-V cells in which ETV2 was silenced by siRNA. ELT3-V cells transfected with ETV2 or Scr siRNA were exposed to arsenite (0.5 mM) for 40 minutes and SG formation was quantified by immunofluorescence using the marker G3BP1 (Anderson & Kedersha, 2002; Kosmas *et al*, 2021). ELT3-V cells transfected with Etv2 siRNA had a significantly increased number of SGs after arsenite treatment compared to cells transfected with SCR siRNA (1.5-fold increase; Figure 3K-L). These data suggest that ETV2 plays a role in SG assembly in response to oxidant stress.

### Syk inhibition regulates Parpbp expression

An examination of the PANDA analysis showed that ETV2 regulated 116 unique genes in the SykI network and 117 unique genes in the rapamycin network. Parp1 Binding Protein (PARPBP) is an important component of DNA replication and damage response pathways and is differentially expressed in various cancers (Feng *et al*, 2014; Uhlen *et al*, 2015; Varisli, 2013). Based on our microarray data and PANDA analysis, we found that Parpbp was regulated by Etv2 uniquely under SykI treatment and demonstrated the highest fold change between treatments compared to other ETV2-regulated genes within that network (Table 2).

**Table 2.**
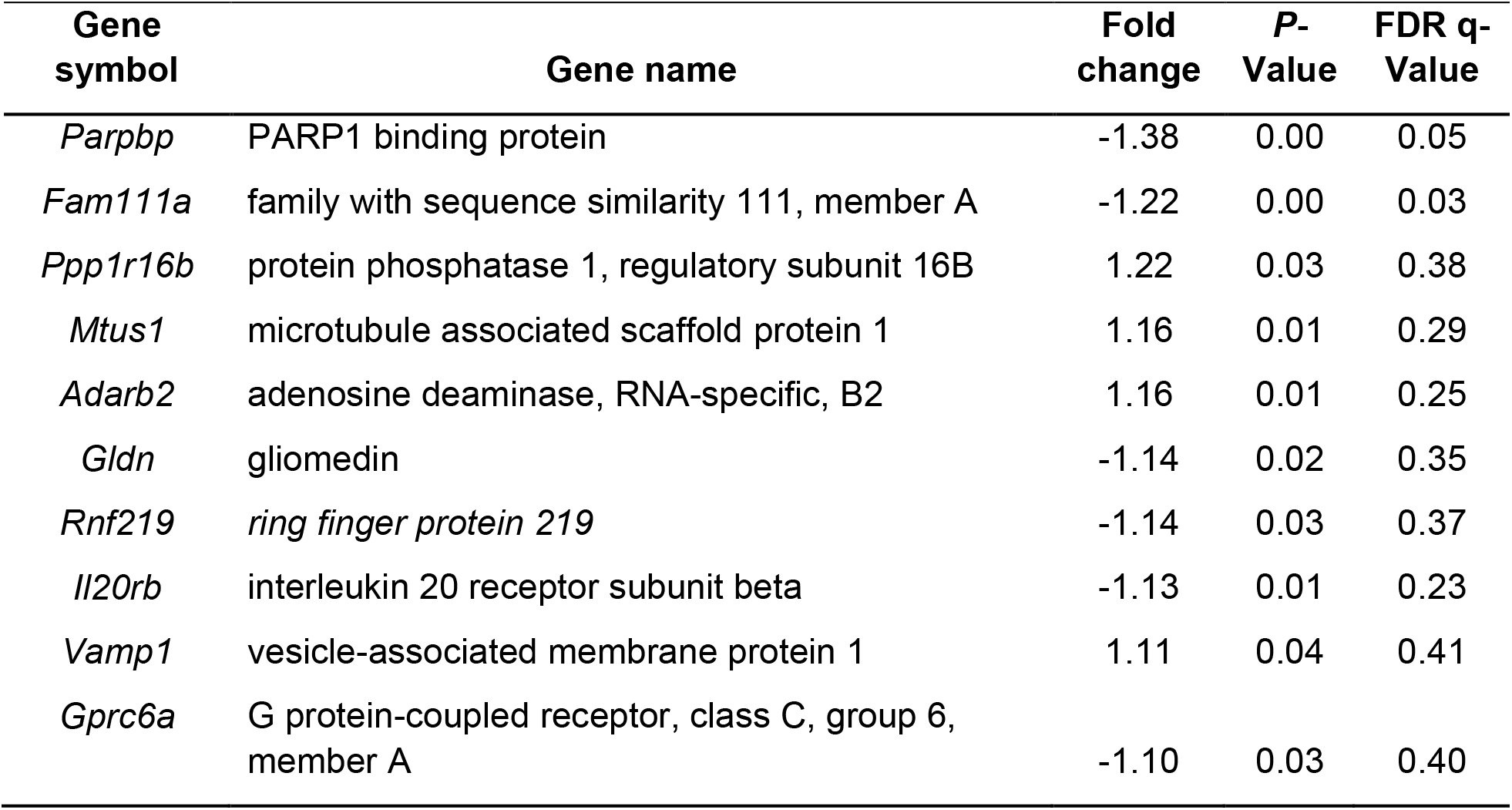
List of top 10 genes identified with PANDA analysis regulated by *Etv2* included in the SykI-specific network.

RT-qPCR analysis showed that both treatments significantly reduced Parpbp expression compared to DMSO, and the reduction was of significantly greater magnitude in the rapamycin treatment group compared to SykI (Figure S2A). We hypothesized that the relatively higher levels of Parpbp mRNA with SykI treatment compared to rapamycin treatment are due to ETV2 nuclear translocation. Accordingly, silencing of ETV2 reduced overall Parpbp expression (Supplementary Figure 2B), as well as abrogated the difference in Parpbp expression level between the two treatment groups (Figure 4A), thus confirming the important role for ETV2 in Parpbp regulation in Tsc2-deficient cells.

**Figure 4.**
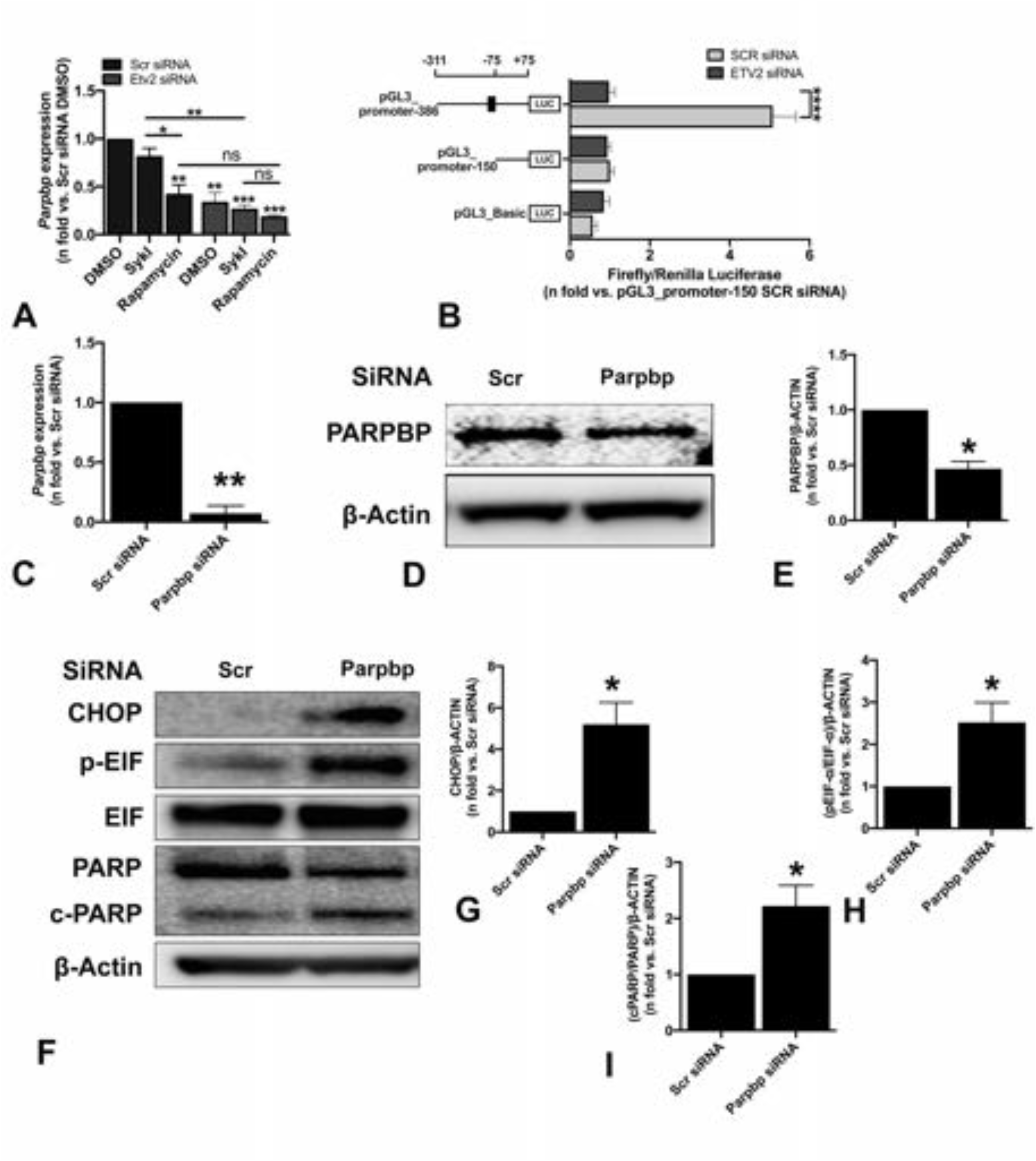
ETV2 silencing-induced changes in PARPBP leads to ER stress and cell death in Tsc2-deficient cells. A) ELT3-V cells were transfected with Scr siRNA and Etv2 siRNA for 24hr and treated with DMSO, SykI, or rapamycin for an additional 24hr. RT-qPCR was carried out on an equal amount of total RNA (1 µg) converted to cDNA to analyze the transcript levels of *Parpbp* gene. The histogram represents fold change in mRNA expression relative to Scr siRNA with DMSO treatment. Data are means ± SEM of at least three independent experiments (* P < 0.05, **P < 0.01, ***P < 0.001; 2-Way Anova Tukey Multiple Comparison). B) ELT3-V cells were transfected with Scr siRNA and Etv2 siRNA for 24hr followed by transfection with internal control, pGL3_Renilla plasmid, and pGL3_Basic vector or PARPBP promoter constructs for an additional 24hr. Firefly luciferase and renilla luciferase activities of negative control, pGL3_Basic vector and PARPBP promoter constructs with and without ETV2-binding site, pGL3_promoter-386 and pGL3_promoter-150 were measured. Each set of luciferase data is normalized to internal renilla control. Data are presented relative to the pGL3_promoter-150 Scr siRNA sample. Data are means ± SEM of at least three independent experiments (****P < 0.0001; Student’s t-test). C) ELT3-V were transfected with either Scr or Parpbp siRNA for 48h. Real-time qPCR analysis of rat Parpbp mRNA was performed. The histogram represents fold change in mRNA expression compared to SCR control. Data are means ± SEM of at least three independent experiments (**P < 0.01; Student’s t-test). D-E) Equal amounts of protein from whole-cell lysates of ELT3-V transfected with either Scr or Parpbp siRNA for 48h were analyzed by Western blotting with antibodies against Parpbp and β-actin. The ratio of Parpbp to β-Actin density was expressed as the fold-change relative to scrambled siRNA. Data are means ± SEM of at least three independent experiments (*P < 0.05; Student’s t-test). F-I) Equal amounts of proteins were separated by electrophoresis and transferred to a PVDF membrane which were reacted with antibodies against CHOP, pEIF-α, total EIF-α, cleaved (c)-PARP, and total PARP. β-Actin was used as a loading control. Ratios of pEIF-α/EIF-α (E), CHOP (F), and c-PARP/PARP(G) to β-Actin density was expressed as fold change to Scr siRNA. Data represent means ± SEM of at least three independent experiments (*P < 0.05, Student’s t-test). **Figure 4-Source data 1** Table showing data from experiments plotted in Figure 4A, 4B, 4C, 4E, 4G, 4H, and 4I. **Figure 4-Source data 2** Original, unedited western blot images and original images with each band labeled for Figure 4D **Figure 4-Source data 3** Original, unedited western blot image files and original images with each band labeled for Figure 4F.

Genome-wide analysis has revealed that ETV2 is an ETS factor with various transcriptional targets containing the consensus sequence of 5’-CCGGAA/T-3’ and a core GGA/T motif (Lee *et al*, 2019; Wei *et al*., 2010). To examine whether ETV2 transcriptionally regulates Parpbp, Parpbp promoter fragments with one or no ETV2 consensus binding sequence were cloned into pGL3_Basic luciferase vector and transfected separately into ELT3-V cells with or without ETV2 silencing to determine promoter activity. The ELT3-V cells without ETV2 silencing transfected with pGL3_promoter-386 construct (i.e., the −311 to +75 bp putative promoter region) with one ETV2 consensus binding site displayed significantly higher luciferase activity compared to cells transfected with the pGL3_promoter-150 construct (i.e., the −75 to +75 bp putative promoter region) with no ETV2 consensus binding site (Figure4B). Additionally, silencing of ETV2 significantly reduced the luciferase activity of the pGL3_promoter-386 construct but showed no effect in the pGL3_promoter-150 construct. These results demonstrated the role of ETV2 in Parpbp transcriptional regulation.

### PARPBP silencing leads to ER stress in Tsc2-deficient cells

PARPBP silencing has been implicated in increased apoptosis and ER stress in myeloid leukemia cells (Nicolae *et al*., 2019). We hypothesize that ETV2-dependent ER stress and increased cell death in Tsc2-deficient cells is mediated by changes in PARPBP expression. To specifically investigate the role of Parpbp in the induction of ER stress and cell death in Tsc2-deficient cells, we silenced PARPBP in ELT3-V cells with Parpbp siRNA. Parpbp siRNA resulted in approximately 80% and 50% reduction in Parpbp mRNA and PARPBP protein, respectively compared to Scr siRNA (Figure 4C-E). Consequently, there was a significant increase in ER stress markers, including CHOP and p-EIF-α (Figure 4F-H), and a significant increase in cell death marker, cleaved-PARP (Figure 4I). These observations support our hypothesis that the regulation of Parpbp expression by ETV2 contributes to ER stress and increased cell death in Tsc2-deficient cells following ETV2 silencing.

### ETV2 silencing leads to Tsc2-deficient cell death *in vivo*

To determine the effect of ETV2 silencing on Tsc2-deficient cell survival *in vivo*, ELT3-V-Luciferase cells were transiently transfected with SCR or Etv2 siRNA for 24 hours. Cells were intravenously injected into female SCID mice and the level of bioluminescence was evaluated in the lungs 4 hours post-injection. Similar levels of bioluminescence intensity were observed in mice injected with both Scr siRNA or Etv2 siRNA transfected cells (Figure 5A-B). By 24H and 48H post-injection, there was a decrease in bioluminescence in both groups; however, the bioluminescence intensities in the mice injected with ETV2-silenced cells were significantly lower compared to mice injected with Scr siRNA transfected cells (Figure 5A-B). We then measured rat-specific DNA in cDNA samples obtained from lung tissue and peripheral blood using RT-qPCR. The levels of rat-specific DNA in both lung tissue and peripheral blood were significantly lower in the mice injected with ETV2 silenced cells compared to SCR control (Figure 5C-D). Consistent with our *in vitro* findings, these data suggest that ETV2 silencing leads to Tsc2-deficient cell death *in vivo*.

**Figure 5.**
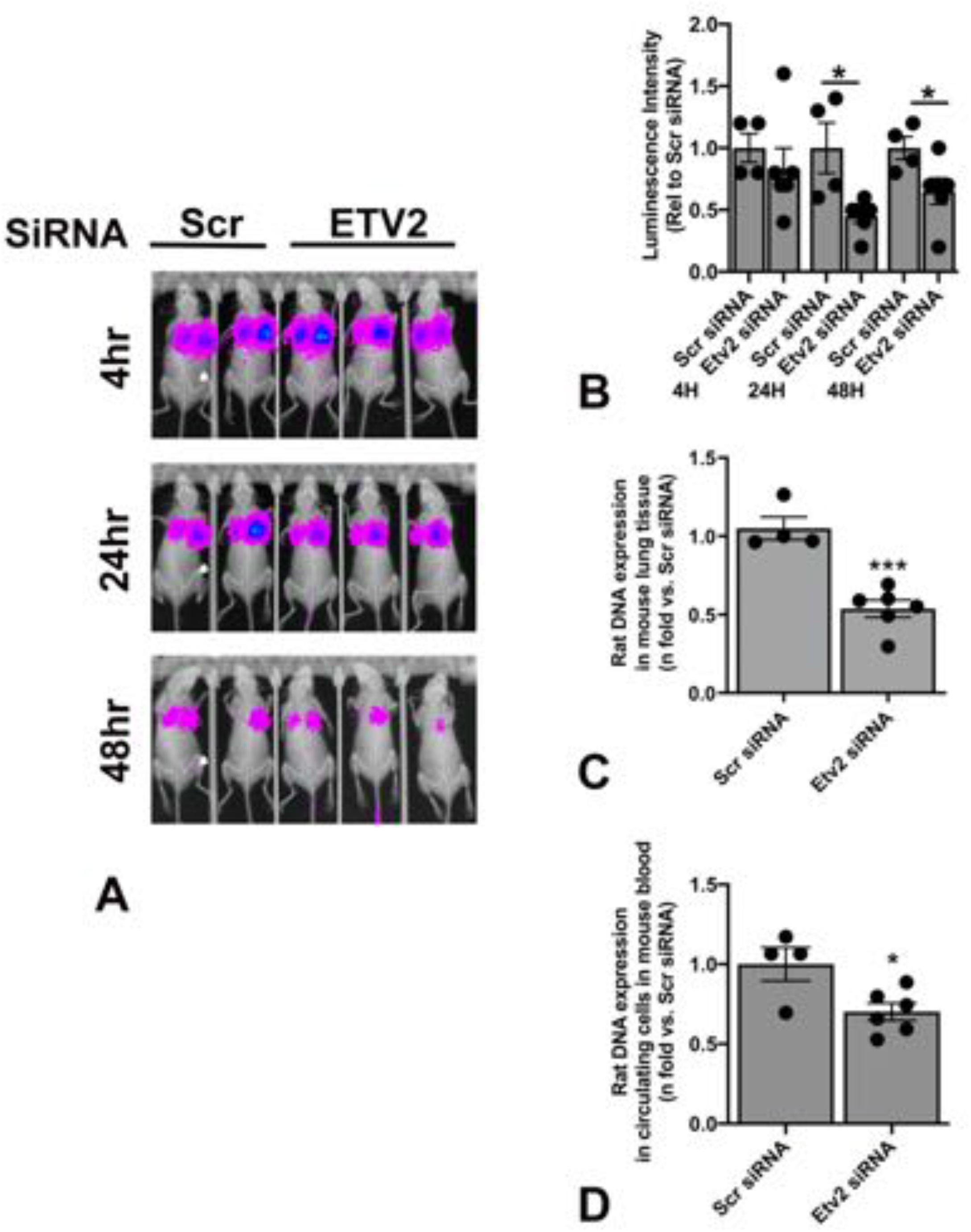
Silencing ETV2 leads to Tsc2-deficient cell death *in vivo*. A) ELT3-V-Luciferase cells were transfected with Scr and Etv2 siRNA for 24hr. 1×10^6^ cells were injected in C.B17 Scid mice (Scr siRNA n=4; Etv2 siRNA n=6) via lateral tail vein injection. Bioluminescence was measured using Bruker In-Vivo Xtreme to show lung colonization of ELT3-V-Luciferase cells at 4hr, 24hr, and 48hr. Representative images are presented. B) Net luminescence intensity was accessed for each mouse at each time point. Histogram reveals a decrease in luminescence intensity in the Etv2 siRNA group relative to the Scr siRNA group at each time point. (*P < 0.05; Student’s unpaired t-test). C) 48hr post-injection, aortic blood was collected from each mouse. Levels of circulating cells from Scr siRNA (n = 4) and Etv2 siRNA (n = 6) mice were measured by RT-qPCR using rat-specific primers. The histogram represents fold change in rat DNA expression relative to Scr siRNA. D) Rat DNA in the mouse lungs harvested 48hr post-injection from Scr siRNA (n = 4) and Etv2 siRNA (n = 6) mice were measured by RT-qPCR using rat-specific primers. The histogram represents fold change in rat DNA expression relative to Scr siRNA. **Figure 5-Source data 1** Table showing data from experiments plotted in Figure 5B, 5C and 5D.

### ETV2 is expressed in human LAM samples

Analysis of single-cell RNA-seq data from 3 LAM lung samples using Seurat identified 7 different unique clusters of cells, including alveolar type II (AT2) cells, conventional dendritic cells (cDC), endothelial cells, fibroblasts, macrophages, natural killer cells, and LAMCORE (Guo *et al*., 2020). A total of 121 LAMCORE cells defined as those which expressed (>0) LAM markers, PMEL, FIGF, ACTA2, and VEGFD(Guo *et al*., 2020), were identified (Figure 6A). One hundred LAMCORE cells were clustered distinctly, 13 were clustered within fibroblast population, and 9 LAMCORE cells were distributed within other cell populations. We also examined ETV2 expression (>0) in LAMCORE cells and demonstrated that 96% of LAMCORE cells were positive for ETV2 expression (Figure 6A).

**Figure 6:**
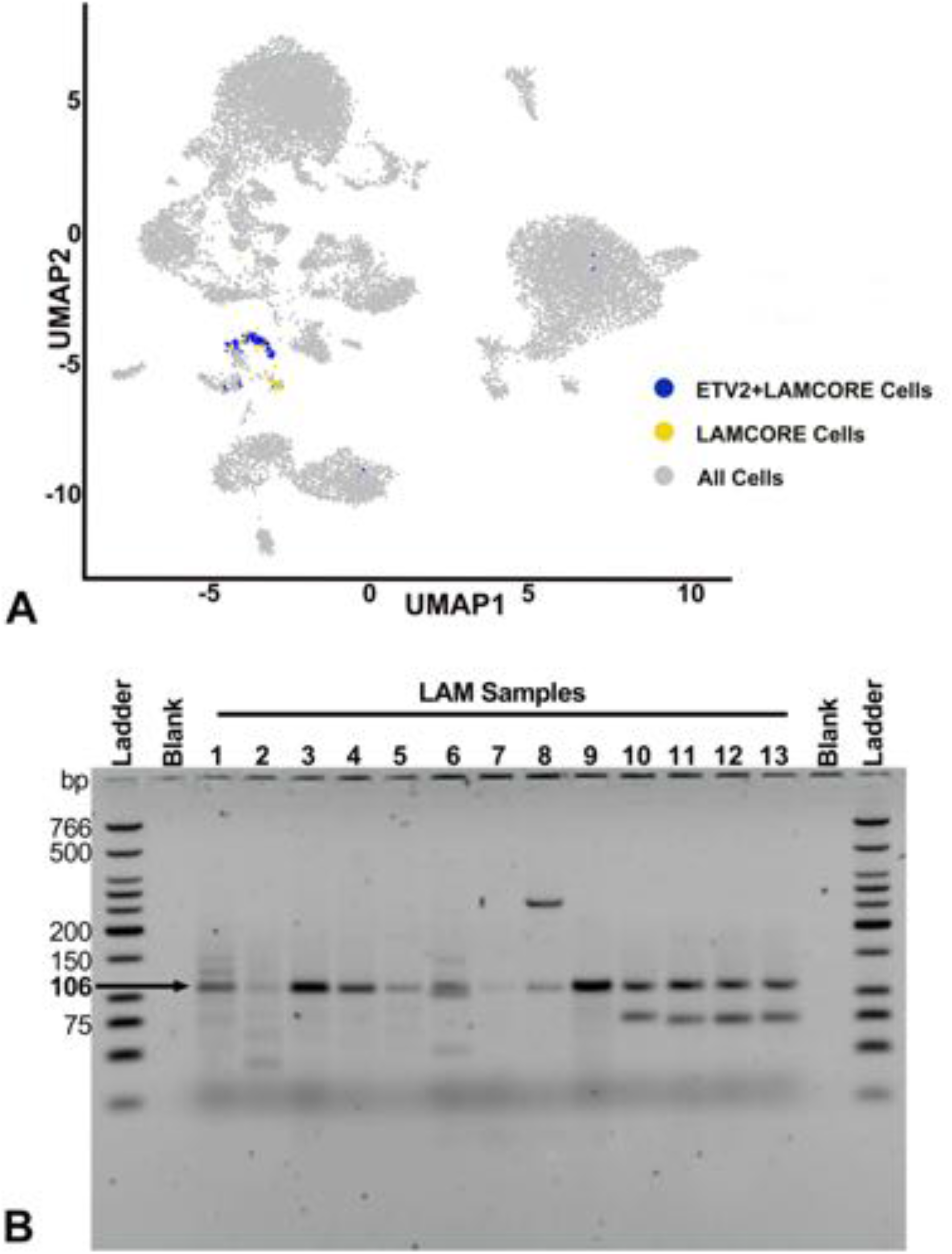
ETV2 expression in human LAM samples. A) Single cells RNAseq analysis of LAM samples detecting ETV2 expressing cells in the LAMCORE population. Feature plot indicates 15 different clusters of cells (gray) and a distinct cluster of LAMCORE cells (yellow), in the LAM samples. ETV2-expressing cells within the LAMCORE cells are indicated in blue. B) Agarose gel electrophoresis analysis of ETV2 expression in cultured LAM cells isolated from explanted lungs of patients undergoing lung transplantation (Samples 1-5) and circulating cells from human blood (Samples 6-13). Amplicons of 106 bp were obtained in all samples. PCR product without RNA to ensure no qRT-PCR contamination is indicated by a blank.

Likewise, sorted cultured cells expressing CD44 and CD44v6 (known to have a loss of heterozygosity for TSC2) isolated from 5 different explanted lung samples (Sample 1-5, Figure 6B) and cells expressing CD235a isolated from 8 blood samples from patients undergoing lung transplantation (Sample 6-13, Figure 6B) were analyzed by RT-PCR for ETV2 expression. Primers designed for PCR targeted the amplification of a 106 bp ETV2 mRNA sequence. The PCR product of the target region from each sample was visualized by electrophoresis on 2% agarose gels. All samples yielded a band at 106 bp, demonstrating the expression of ETV2 in all human LAM samples.

## Discussion

Due to the constitutive activation of mTORC1 signaling, the use of mTOR inhibitors such as rapamycin has remained the primary therapeutic approach in the treatment of LAM (Johnson *et al*., 2016; McCormack *et al*., 2011). However, because of the cytostatic effects of rapamycin in Tsc2-deficient cells, there is a need for additional therapeutic targets. Identifying these targets requires a better understanding of death pathways in *Tsc2*-deficient cells. We have previously shown that Syk inhibition has effects both upstream and downstream of mTORC1, and causes decreased proliferation of *Tsc2*-deficient cells *in vitro* and *in vivo*, and similar to mTORC1 inhibition did not lead to cell death (Cui *et al*., 2017). Additionally, a Syk inhibitor, such as R406 was shown to have secondary targets besides Syk kinase (Cui *et al*., 2017; Rolf *et al*., 2015), which can be explored to better understand better cell death pathways in LAM cells. Therefore, we utilized gene profiling and network-based approaches with PANDA analyses to identify potential regulatory pathway(s) that are unique to SykI treatment. These analyses identified a transcription factor, ETV2 regulating gene expressions of various genes uniquely during Syk-I treatment but not with rapamycin treatment.

ETV2 is a well-known regulator of blood and endothelial cell lineages during development (Garry, 2016; Li & Sidell, 2005). To date, the role of ETV2 in *Tsc2*-deficient tumors has not been investigated. Our data demonstrated that although Syk-inhibition does not affect expression of ETV2 at mRNA or protein levels, it prompted the translocation of ETV2 into *Tsc2*-deficient cell nuclei. Importantly, rapamycin treatment does not affect the expression or nuclear localization of ETV2, suggesting that ETV2 may drive a potential mTORC1-independent transcriptional pathway in *Tsc2*-deficient cells. The presence of cAMP response element (CRE) sequences and therefore regulation of ETV2 by PKA signaling has been shown previously (Yamamizu *et al*, 2012). Likewise, the interaction between Syk and PKA has also been described (Yu *et al*, 2013). Therefore, the nuclear translocation of ETV2 during Syk inhibitor treatment could potentially be driven in a PKA-dependent manner. Further elucidation of this pathway will require additional investigation.

Known target genes of ETV2, acting as a transcription factor, are genes involved in the regulatory networks for hematopoietic and endothelial lineages, including Flk1 (Kim *et al*, 2019). We used PANDA network analysis and identified multiple genes that could be potential transcriptional targets of ETV2 following Syk-inhibition in Tsc2-deficient cells. The most differentially regulated gene was *Parpbp*. Interaction between ETV1, EWS-ERG fusion genes, and EWS-FLI1 fusion genes, all of which are members of ETS family transcription factors, and PARP1, a PARPBP-interacting partner has been demonstrated in Ewing sarcoma and prostate cancers (Feng *et al*., 2014). Our study is the first to demonstrate an interaction between ETV2 and PARPBP. We showed that SykI treatment resulted in the level of PARPBP expression that was lower than DMSO treatment but significantly higher than rapamycin treatment. SykI treatment-induced nuclear translocation of ETV2, which we also showed to transcriptionally activate PARPBP expression, is likely to be increasing PARPBP expression, negating some of the effects of mTORC1 inhibition.

More importantly, to investigate the functional role of ETV2 in *Tsc2*-deficient cells, we silenced ETV2 and demonstrated that silencing of ETV2 induced ER-stress, leading to increased cell death both *in vitro* and *in vivo*. We also showed that ETV2-dependent cell death is driven by the ER-stress pathway. Our finding of ER-stress-mediated *Tsc2*-deficient cell death extends prior knowledge on the contribution of ETV2 to cell survival and apoptosis (Abedin *et al*., 2014; Singh *et al*., 2019) as well as the susceptibility of *Tsc2*-deficient cells to ER-stress (Kang *et al*., 2011; Ozcan *et al*., 2008). It was recently shown that TSC2 physically interacts with the stress granule protein G3BP1, and that stress granules are increased in *Tsc2*-deficient cells (Kosmas *et al*., 2021). Stress granules may represent a mechanism to temporarily sequester transcripts during transient stress, including oxidant stress. Our data showed that silencing ETV2 increased stress granules in *Tsc2*-deficient cells in the setting of short term arsenite-induced oxidant stress, suggesting that multiple mechanisms impact stress granules in TSC. Furthermore, we also demonstrated that PARPBP silencing, similar to silencing of ETV2, resulted in increased ER stress and increased cell death in *Tsc2*-deficient cells. This suggested that ETV2 induces ER stress and increased cell death in *Tsc2*-deficient cells via PARPBP regulation (Figure 7).

**Figure 7:**
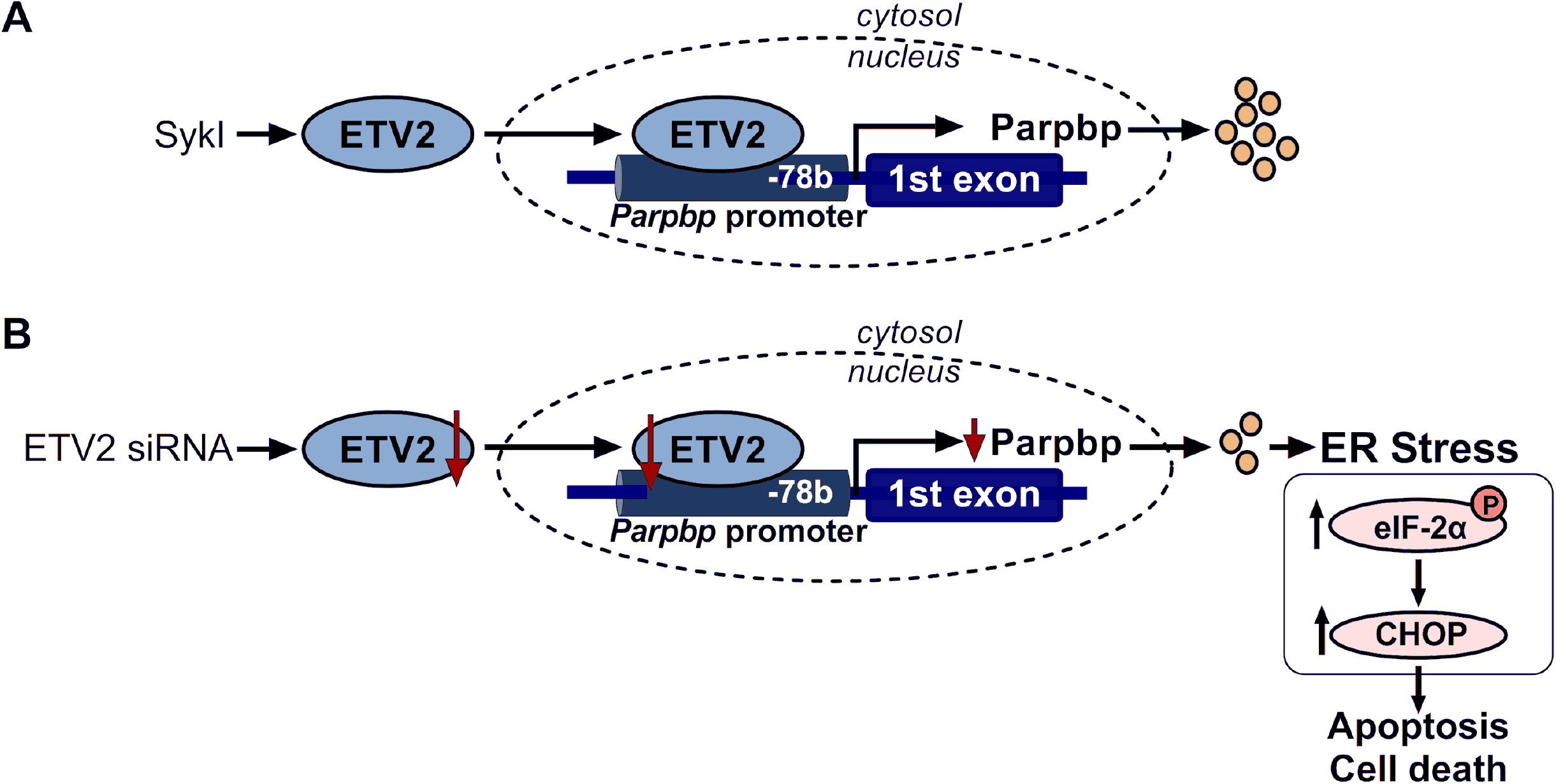
Schematic representation of ETV2 nuclear translocation and PARPBP regulation. A) In Tsc2-deficient cells, Syk inhibition (SykI) drives ETV2 to the nucleus, where it binds to its consensus sequence in the PARPBP promoter at position −78 base pair to transcriptionally activation of *Parpbp* gene expression. B) In Tsc2-deficient cells, decreased ETV2 expression results in reduced PARPBP mRNA and protein. Reduction in ETV2 or PARPBP protein levels induces ER stress, with increased phosphorylation of eIF-2a and CHOP expression, resulting in increased apoptosis and cell death.

ELT3 cells share some important common features with LAM cells including tuberin deficiency, activation of mTOR, and uterine origin (Guo *et al*., 2020). We elected to use ELT3 cells in an *in vivo* model of *Tsc2*-deficient cell homing to the lung, where we again showed that ETV2 deficiency leads to significantly decreased cell survival in the lung. Finally, we validated ETV2 expression in human LAM cells in three different ways. First, we identified LAM cells in the recently published single-cell RNAseq data and found that these cells overwhelmingly expressed ETV2. Second, in a mixed cell population isolated from LAM lungs, we found that cells with loss of heterozygosity for *TSC2* expressed ETV2. Finally, LAM cells isolated from peripheral blood from female patients with LAM also expressed ETV2.

Taken together, the combination of the *in vitro, in vivo* and human data strongly support our hypothesis of a critical role for ETV2 in *Tsc2*-deficient cell survival. Therefore, identifying molecules that target ETV2 expression or its nuclear trafficking could potentially be of important therapeutic benefit in LAM and other tumors driven by mTORC1 activation.

## Supplementary Figures

**Supplementary Figure 1:**
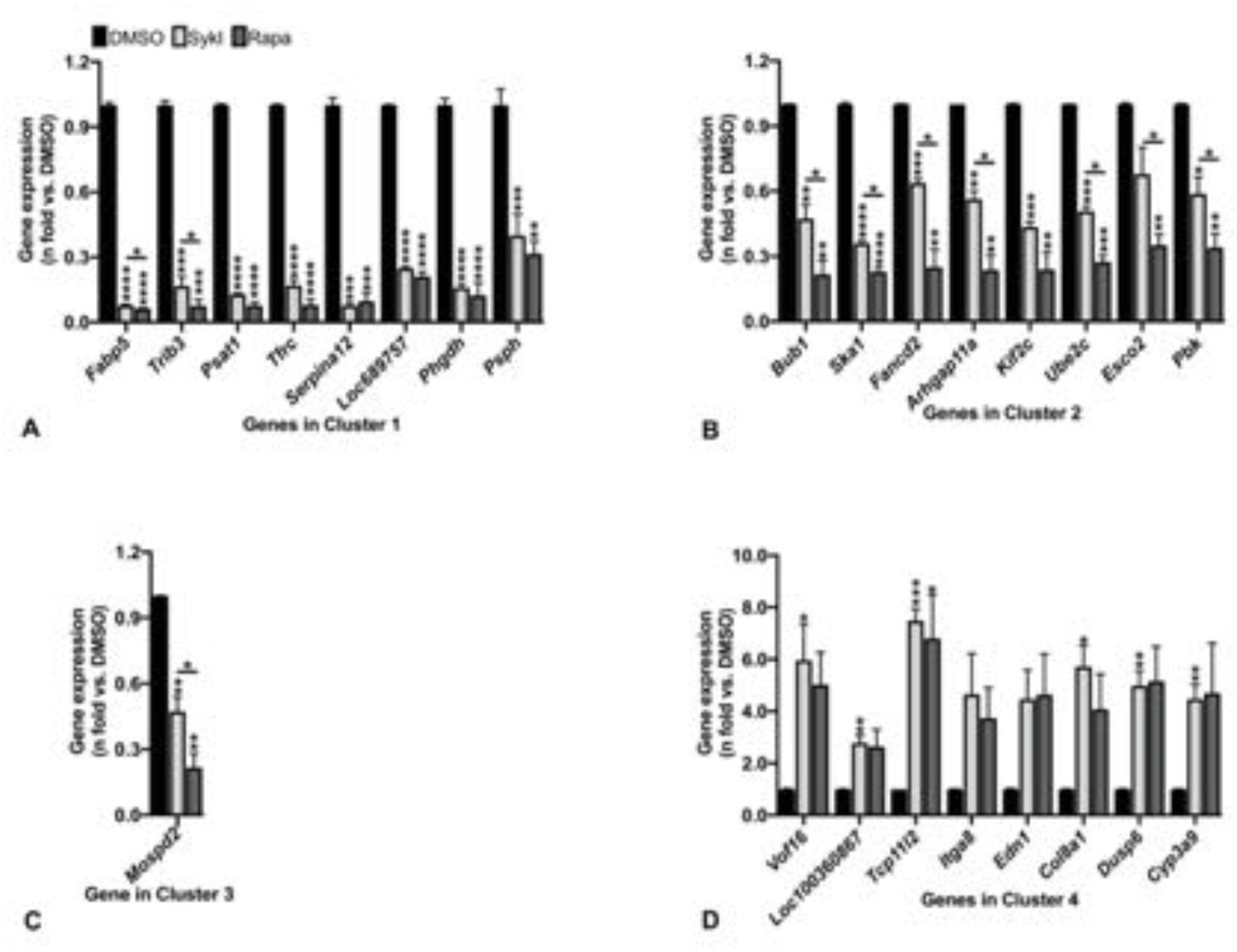
Real-time qPCR validation of microarray gene expression data. A-D. RT-qPCR was performed for microarray gene expression data for topmost differentially regulated genes from (A) Cluster 1, (B) Cluster 2, (C) Cluster 3, and (D) Cluster 4 in the heatmap (Figure 1A). Data are means ± standard error of mean (SEM) of four independent experiments (*P < 0.05*, **P < 0.01, ***P < 0.001, ****P < 0.0001; 2-Way Anova Tukey’s multiple comparison test). **Supp Figure 1-Source data 1** Table showing data from experiments plotted in Supplementary Figure 1A, 1B, 1C and 1D.

**Supplemental Figure 2:**
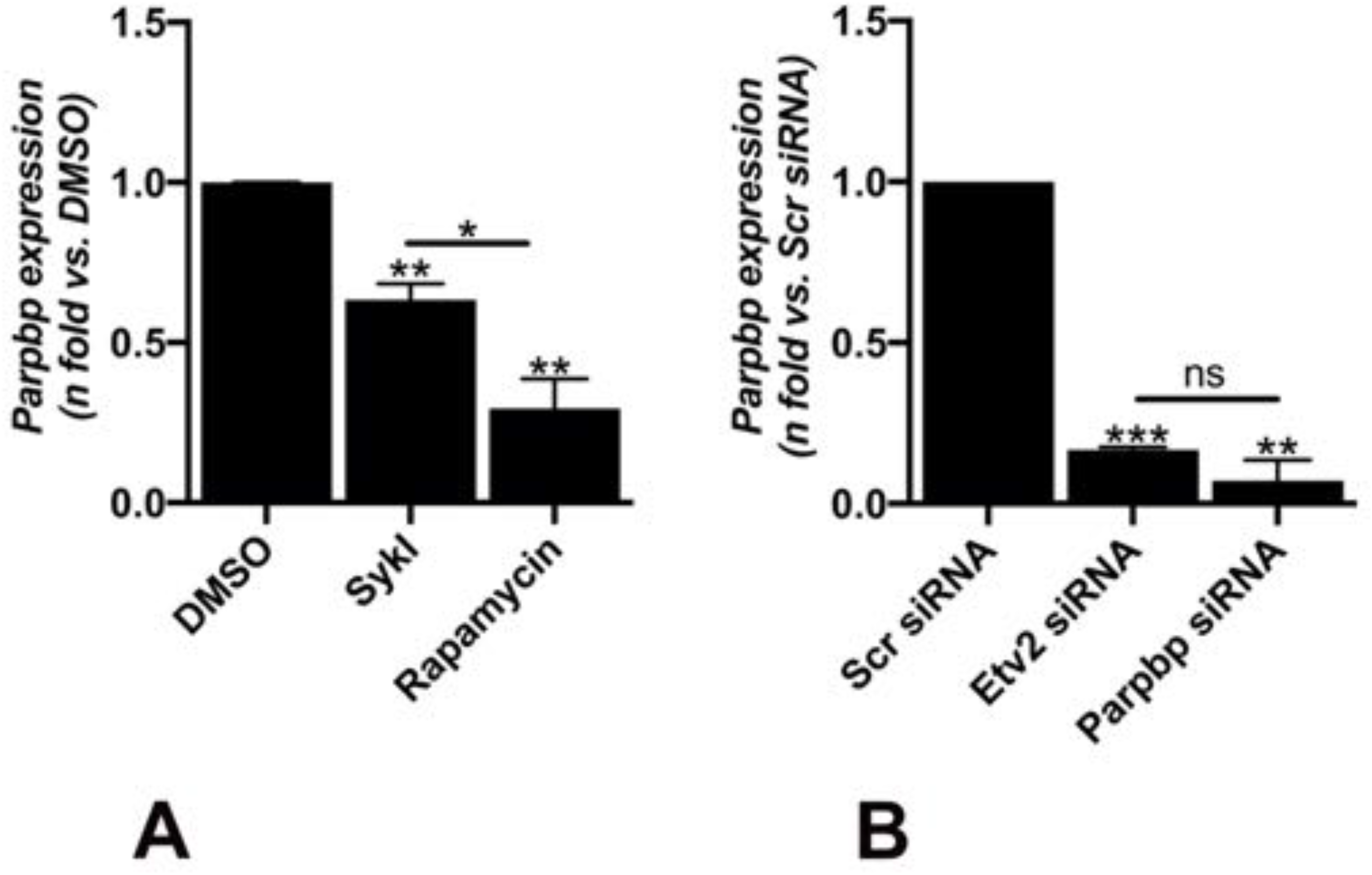
Syk inhibition and ETV2 silencing reduce *Parpbp* expression. A) Real-time qPCR analysis of rat *Parpbp* mRNA in *Tsc2*-deficient (ELT3-V) cells treated with SykI, or rapamycin for 24H compared to DMSO-treated cells. Data represent mean ± SEM of three independent experiments. B) Real-time qPCR analysis of rat *Parpbp* mRNA in ELT3V cells transfected with Etv2 siRNA and Parpbp siRNA for 48hr. Data represent mean ± SEM of three independent experiments. **Supp Figure 2-Source data 1** Table showing data from experiments plotted in Supplementary Figure 2A and 2B.

## Notes

**Funding:** This work was supported in part by National Institutes of Health (U01-HL 131022 to S. El-Chemaly and T32HL007633-35 to J. Ng) and the Division of Intramural Research, National Institutes of Health, National Heart, Lung, and Blood Institute and the Anne Levine LAM Research Fund (S. El-Chemaly)

### Competing Interest Statement

The authors have declared no competing interest.

